# Deep learning-based cytoskeleton segmentation for accurate high-throughput measurement of cytoskeleton density

**DOI:** 10.1101/2024.05.27.596126

**Authors:** Ryota Horiuchi, Asuka Kamimura, Yuga Hanaki, Hikari Matsumoto, Minako Ueda, Takumi Higaki

**Author notes:** **Corresponding author:** T. Higaki.

## Abstract

Microscopic analyses of cytoskeleton organization are crucial for understanding various cellular activities, including cell proliferation and environmental responses in plants. Traditionally, assessments of cytoskeleton dynamics have been qualitative, relying on microscopy-assisted visual inspection. However, the transition to quantitative digital microscopy has introduced new technical challenges, with segmentation of cytoskeleton structures proving particularly demanding. In this study, we examined the utility of a deep learning-based segmentation method for accurate quantitative evaluation of cytoskeleton organization using confocal microscopic images of the cortical microtubules in tobacco BY-2 cells. The results showed that, although conventional methods sufficed for measurement of cytoskeleton angles and parallelness, the deep learning-based method significantly improved the accuracy of density measurements. To assess the versatility of the method, we extended our analysis to physiologically significant models in the context of changes in cytoskeleton density, namely *Arabidopsis thaliana* guard cells and zygotes. The deep learning-based method successfully improved the accuracy of cytoskeleton density measurements for quantitative evaluations of physiological changes in both stomatal movement in guard cells and intracellular polarization in elongating zygotes, confirming its utility in these applications. The results demonstrate the effectiveness of deep learning-based segmentation in providing precise and high-throughput measurements of cytoskeleton density, and has the potential to automate and expedite analyses of large-scale image datasets.

## Introduction

Cytoskeleton organization in plant cells undergoes dynamic changes in response to internal factors, such as cell-cycle progression, and external stimuli, including environmental stress, playing a crucial role in various cellular functions. For instance, immediately following mitosis, a phragmoplast containing microtubules and actin filaments is formed during cytokinesis and then quickly disappears after the completion of cell division (Maeda et al. 2020; Schmidt-Marcec et al. 2024). Subsequently, plant cells establish cortical microtubules anchored to the plasma membrane, which are crucial for directing the orientation of cellulose microfibrils that determine the direction of cell elongation (Kumagai et al. 2001; Paredez et al. 2006; Lucas 2021). In addition, plant cells can rapidly degrade these cortical microtubules in response to hyperosmotic stress (Fujita et al. 2013; Dou et al. 2018). As these examples illustrate, understanding how cytoskeleton organization functions and contributes to cell proliferation and environmental responses, together with the molecular mechanisms that regulate these processes, represents a critical research topic in plant cell biology. Microscopic observation of the cytoskeleton is indispensable to address these research objectives. Traditionally, cytoskeleton organization was qualitatively evaluated through visual inspection. However, in recent years, quantitative analysis of cytoskeleton organization through digital microscopic image analysis has become the standard analytical approach (Paez-Garcia et al. 2018).

In microscopic image analysis, the process of identifying specific cell structures within the images is termed segmentation (Legland et al. 2016; Laan et al. 2023). This procedure is fundamental and critical to analyze the cytoskeleton and organelles. In images with fluorescently labeled cytoskeletons, cytoskeletal regions are typically defined based on fluorescence intensity, where regions with intensity above a predetermined threshold are identified as cytoskeleton (Higaki et al. 2010; Li et al. 2022; Hembrow et al. 2023). Two primary methods are used to determine this threshold. One method involves the researcher manually setting the intensity that distinguishes the cytoskeleton from the background for each microscopic image. Although this method can yield highly accurate thresholds, it becomes overly labor-intensive when numerous images must be analyzed and it may suffer from a lack of reproducibility. The alternative method involves automatically setting the threshold using an intensity thresholding algorithm, such as Otsu’s method (Otsu 1979). This method significantly reduces the manual intervention required and ensures reproducibility, although it may occasionally result in decreased accuracy depending on the image properties. In practice, researchers often manually set thresholds to prioritize accuracy. Although both approaches have their advantages and disadvantages, to address plant cell biology problems, manual segmentation has been frequently utilized because the accuracy of the analysis is more important than the human cost (Higaki et al. 2010; Dou et al. 2018). However, manual analysis of temporal changes in cytoskeleton density across multiple samples in time-lapse imagery is impractical, presenting a substantial technical challenge.

To address this challenge, there is a growing need for segmentation methods that are not only efficient but also accurate. Although examples of research on cytoskeleton segmentation are limited, reports of automated segmentation methods that mimic human judgment based on deep learning techniques have emerged mainly in animal cell biology (Özdemir and Reski 2021). Deep learning has been applied to segment microtubules in human HeLa cells (Yue et al. 2023). A pre-trained deep learning model for image segmentation (AIVIA 2-D segmentation) (Kikukawa et al. 2021, 2023) was trained by sets of fluorescence microscopic images of microtubules and corresponding manually thresholded segmentation images from time-lapse data (Yue et al. 2023). The trained model effectively mimicked human visual thresholding, facilitating automated segmentation that closely matches human accuracy. This method not only reduced the workload for researchers, but also ensured consistent and reproducible measurements of microtubule density from time-lapse data (Yue et al. 2023). The application of deep learning in cytoskeleton segmentation has demonstrated promise to enhance accuracy, reproducibility, and throughput. However, systematic comparisons with conventional methods of cytoskeleton segmentation have not yet been conducted, especially in plant cell biology. Understanding the cytoskeleton organization in plant cells is particularly important because of its crucial roles in diverse cellular processes, such as rapid cell growth and response to environmental stimuli. Therefore, the present study aimed to systematically evaluate the effectiveness of a deep learning-based method compared with conventional segmentation methods in analyzing fluorescence microscopic images of actual plant cytoskeletons. By conducting evaluations across three metrics for cytoskeleton organization—angle, parallelness, and density—using an image dataset of cortical microtubules in tobacco BY-2 cells, we sought to determine the optimal and practical approach for plant cytoskeleton segmentation. The results showed that conventional methods were sufficiently accurate for measurement of angle and parallelness; however, a deep learning model-based method, with more than 40 training image sets, was able to measure density with greater precision compared with conventional segmentation methods. Furthermore, the versatility of this method was confirmed for analysis of microtubules and actin filaments in *Arabidopsis thaliana* guard cells, and microtubules in *A. thaliana* zygotes. The method enabled highly precise evaluations of the increase in actin filament density accompanying stomatal closure induced by abscisic acid (ABA), as well as the biased distribution of microtubules during the elongation of zygotes. This study reveals the potential of deep learning-based methods for quantitative evaluation of plant cytoskeleton organization by demonstrating the utility of these techniques in diverse experimental systems.

## Materials and Methods

### Plant materials

To capture confocal microscopic images of cortical microtubules, *Nicotiana tabacum* L. ‘Bright Yellow 2’ (tobacco BY-2) cells stably expressing YFP-tubulin under the control of the cauliflower mosaic virus (CaMV) 35S promoter were utilized (Kojo et al. 2013). The transgenic cells were maintained similarly to wild-type BY-2 cells, diluted 95-fold with modified Linsmaier and Skoog medium supplemented with 2,4-dichlorophenoxyacetic acid at weekly intervals (Kumagai-Sano et al. 2006). The cell suspensions were incubated on a rotary shaker at 130 rpm at 27°C in the dark.

Confocal microscopic images of GFP-mouse talin (mTn) and GFP-tubulin in stomatal guard cells of fully expanded rosette true leaves from 4-to 5-week-old *A. thaliana* plants were obtained from the LIPS database (https://www.higaki-lab.net/lips/) (Higaki et al. 2012, 2013). In addition, to capture confocal images of actin filaments in guard cells treated with ABA, transgenic plants expressing GFP-ABD2 under the control of the CaMV 35S promoter were used (Higaki et al. 2010; Lu et al. 2020). Seeds were sown in soil (Jiffy-7; Sakata Seed Corp., Yokohama, Japan) and grown in a chamber at 23.5°C under a 16-h light/8-h dark cycle, with illumination from an 86.2 μmol m^−2^ s^−1^ light-emitting diode (Plantflec, LH-241PFP-S; NK System, Tokyo, Japan). For ABA treatments, 14-day-old *A. thaliana* seedlings were submerged in a closing buffer [5 mM KCl, 50 μM CaCl_2_, 10 mM MES, pH 6.15 (Tris)] (Sato et al. 2022) containing either 0.1% dimethyl sulfoxide (DMSO) as a control or 10 μM ABA for 5 min. Subsequently, the surfaces of the cotyledons were immediately examined with a confocal microscope.

To obtain two-photon excitation microscopic images of microtubules in *A. thaliana* zygotes, we used plants expressing a microtubule/nucleus marker comprising the 463-bp *EGG CELL1* (*EC1*) promoter, the GFP variant Clover, the TUBULIN ALPHA6 (TUA6), and the NOS terminator in the pMDC99 binary vector (EC1p::Clover-TUA6; coded as MU2228) (Kimata et al. 2016; Curtis and Grossniklaus 2003). Self-pollinated flowers were dissected under a stereomicroscope and the collected ovules were cultivated as previously described (Ueda et al. 2020).

### Microscopy

For confocal imaging of BY-2 cells and *A. thaliana* guard cells, we used a microscope (IX-70; Olympus, Tokyo, Japan) equipped with a CSU-X1 scanning head (Yokogawa, Tokyo, Japan), a 100× objective lens (UPlanSApo, NA = 1.40; Olympus), and a scientific complementary metal oxide semiconductor camera (Prime 95B; Teledyne Photometrics, Tucson, AZ, USA). YFP and GFP were excited with a 488 nm laser and fluorescence was detected through a 510–550 nm band-pass filter.

For two-photon imaging of *A. thaliana* zygotes, we used a laser-scanning inverted microscope (AX; Nikon) equipped with a pulse laser (InSight X3 Dual option; Spectra-Physics). The images were acquired as 31 z stacks with 1-μm intervals using a 40× water-immersion objective lens (CFI Apo LWD WI, NA = 1.15; Nikon) with Immersol W 2010 (Zeiss) immersion medium. Fluorescence signals were detected using the GaAsP PMT detector and a 470/40 nm band-pass filter.

### Image processing and analysis

Image processing was conducted using ImageJ software (Schneider et al. 2012) for conventional techniques, including thresholding and skeletonization of binary images. To enhance cytoskeleton structures in guard cells and zygotes, the FFT Bandpass Filter in ImageJ was applied with Sigma values set to 1.5 and 1.0 pixels as a pre-processing step. Binarization of the cytoskeleton images was performed using Otsu’s thresholding in ImageJ, followed by skeletonization of the binary images using the LPX bilevelThin plugin with default settings (Biel et al. 2022).

Deep learning-based image transformation was conducted using the 2-D segmentation function of the AIVIA image analysis software version 9.8.1 (DRVision, Bellevue, WA, USA) (Kikukawa et al. 2021, 2023). The model is based on the residual channel attention (RCA)-UNet architecture, which enhances the model’s ability to focus on informative features at each resolution level (Ronneberger et al. 2015; Zhang et al. 2018). We used the default settings for model training. A list of the default parameters and their explanations can be found at the following link: https://aivia-software.atlassian.net/wiki/spaces/AW/pages/1797652487/Deep+Learning+Hyperparameters+Settings. The model was trained using raw confocal images and manually segmented binary images as input data in AIVIA. Following the training, it generated segmented images of the cytoskeleton from the input confocal images. The segmented images were subsequently processed in ImageJ, where binarization was applied using Otsu’s thresholding method.

To quantify cytoskeleton organization, three numeric metrics—angle, parallelness, and density— were measured in the skeletonized images using the ImageJ plugin LPX LineFeature (Higaki 2017; Yoshida et al. 2023). Angle represents the average orientation of the cytoskeleton and ranges from 0 to 180 degrees. Parallelness indicates the alignment of cytoskeleton structures, reaching a maximum value of 1 when all structures are aligned in the same direction and approaching a minimum value of 0 as orientations diverge. Density represents the total length of skeletonized cytoskeleton per unit area, providing an estimate of cytoskeleton coverage.

Statistical analyses were conducted using the R statistical analysis software (https://www.r-project.org/). The training and test image sets for cortical microtubules in BY-YTRF cells are publicly accessible on figshare under the CC BY 4.0 license (https://doi.org/10.6084/m9.figshare.27634116.v1). The deep learning models for enhancing cortical microtubule structures, designed for use with AIVIA software, are also available on figshare (https://doi.org/10.6084/m9.figshare.27683220.v1).

## Results

### Deep learning segmentation using cortical microtubule images of tobacco BY-2 cells

To develop a training image dataset for deep learning-based cytoskeleton segmentation, we initially captured 498 confocal images of cortical microtubules labeled with YFP-tubulin in tobacco BY-2 cells (Kojo et al. 2013). These images were randomly divided into 400 training images to train the deep learning segmentation model and 98 test images to evaluate the segmentation accuracy. To create the ground-truth data for cytoskeleton segmentation, these images underwent manual segmentation based on manually determined thresholds (Fig. 1a, Manual segmentation). Subsequently, a size filter was applied to omit globules smaller than 10 pixels (Fig. 1a, Size filtering). Finally, the segmented images were converted into binary format and skeletonized to measure cytoskeleton metrics, comprising angle, parallelness, and density (Fig. 1a, Skeletonization and Measurements). To compare the deep learning-based cytoskeleton segmentation with conventional methods, we modified the image processing pipeline of the ground-truth images by replacing manual segmentation with Otsu’s method, a representative threshold determination algorithm, to measure cytoskeleton metrics (Fig. 1b). We then developed the image processing pipeline as a proposed method: we trained a pre-trained deep learning model for image segmentation (Kikukawa et al. 2021) with raw images and the manually segmented binary images. The training image set was incrementally increased from *N* = 2 to *N* = 400, and the test set of 98 images was used for accuracy assessment. The trained models should emphasize the cytoskeleton structures (Fig. 1c, DL-based image transformation), and the enhanced images were then automatically binarized using Otsu’s method for skeletonization and metric measurements (Fig. 1c).

**Figure 1.**
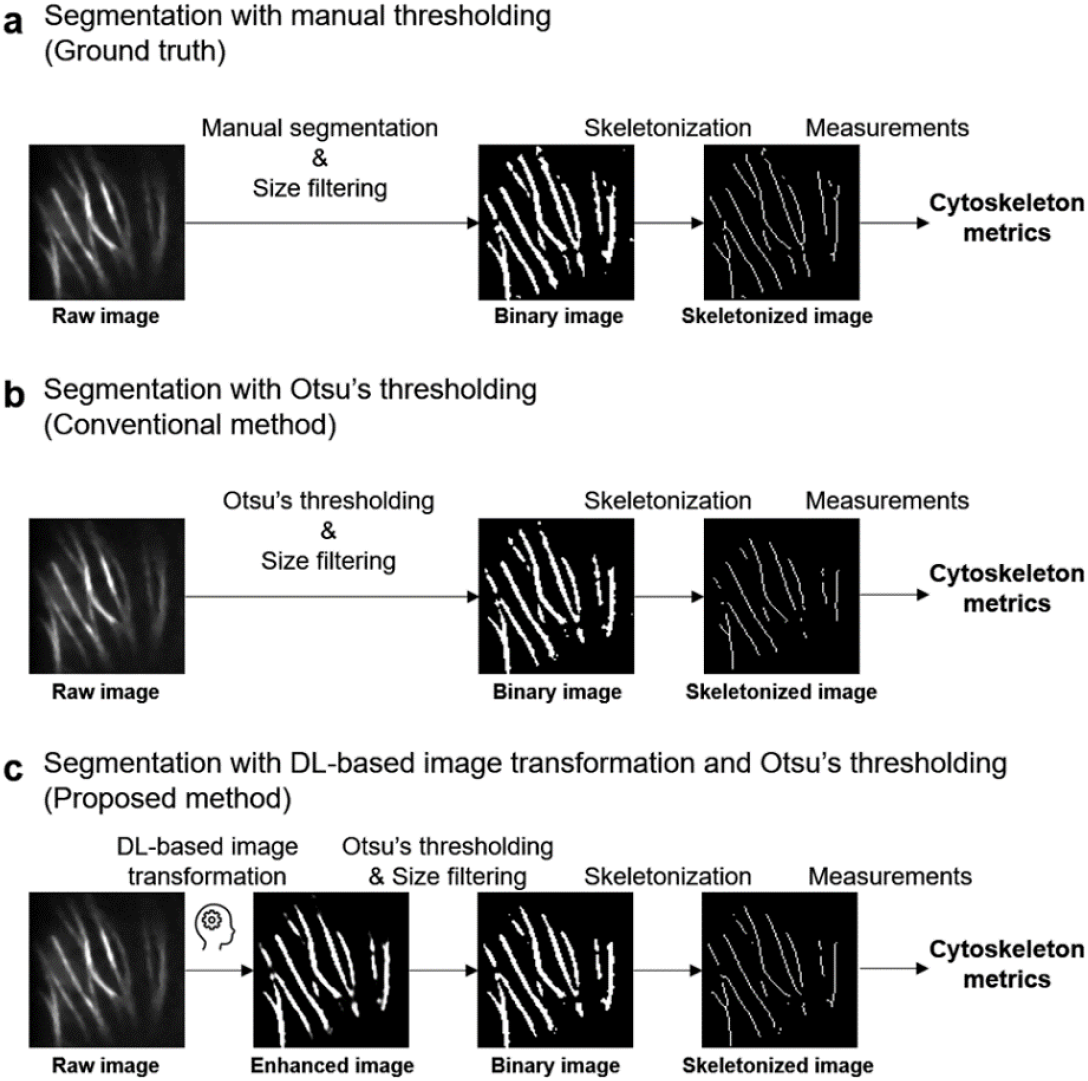
Image processing pipelines for evaluation of the accuracy of the proposed cytoskeleton segmentation method. (a) The ground-truth method begins with manual intensity thresholding of raw confocal images, followed by size filtering to remove globules smaller than 10 pixels, skeletonization, and measurement of cytoskeleton metrics. (b) The conventional method employs Otsu’s method for automated thresholding, followed by binarization, size filtering, skeletonization, and measurements, as for the ground-truth method. (c) The proposed method utilizes a deep learning model trained with raw and manually thresholded binary images for image transformation, enhancing the cytoskeleton structures. The subsequent steps of binarization, skeletonization, and measurements are identical to those in the conventional method. A representative image of cortical microtubules in tobacco BY-2 cells is shown as an example. The image width of the square is 18.5 μm.

From this analysis, we obtained measurements of the angle, parallelness, and density of the cortical microtubules, as representative cytoskeleton metrics (Higaki 2017), for the 98 test images based on ground truthing (Fig. 1a), the conventional method (Fig. 1b), and the proposed method (Fig. 1c). To evaluate the segmentation accuracy of the conventional and proposed methods, we generated scatter plots to compare their metric values with those obtained by ground truthing (Figs. 2–4).

**Figure 2.**
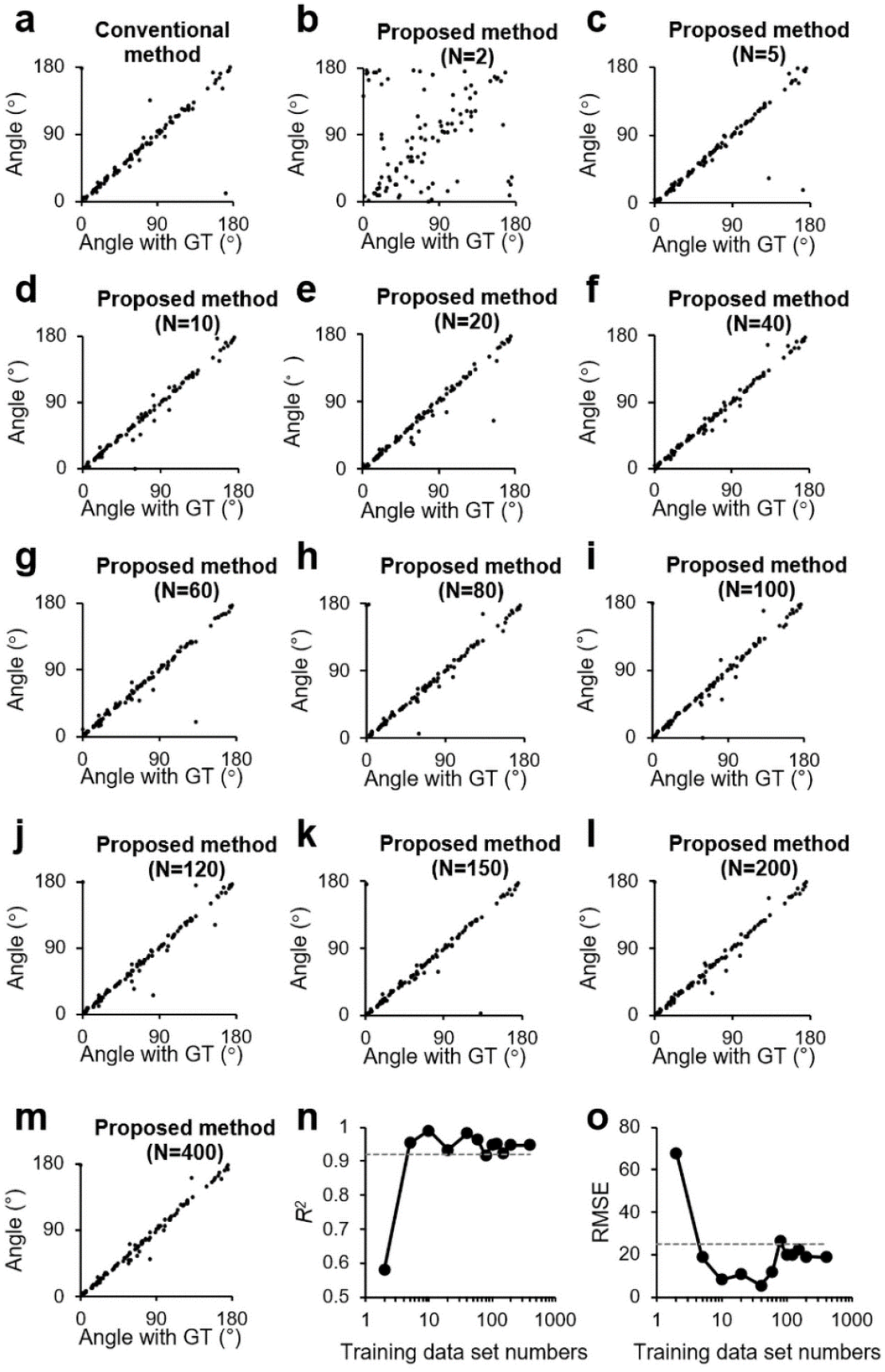
Evaluation of image processing pipelines for measurement of cytoskeleton angles. (a) Scatterplot comparing cytoskeleton angle values measured by the ground truth (GT) against those obtained via the conventional method. (b–m) Scatterplots comparing GT cytoskeleton angle values to those measured by the proposed method, varying the number of training image datasets at *N* = 2 (b), 5 (c), 10 (d), 20 (e), 40 (f), 60 (g), 80 (h), 100 (i), 120 (j), 150 (k), 200 (l), and 400 (m). (n, o) Plot of *R*^2^ (n) and root mean squared error (RMSE) (o) as an accuracy metrics, shown against the number of training datasets in the proposed method. The dashed line represents the value of 0.921 and 24.9 for the conventional method, respectively.

The angle represents the average orientation of the cytoskeleton in the images (Higaki 2017), which varied from 0 to 180 degrees in the test images. Comparison of the ground-truth and conventional method measurements indicated they were well aligned (Fig. 2a). To quantitatively assess the accuracy of segmentation and density measurements, we used the coefficient of determination (*R*^2^) and the root mean squared error (RMSE) to evaluate the discrepancies between the measured values obtained by each method and the ground truth. A higher *R*^2^ value indicates a stronger correlation with the ground truth, while a lower RMSE value indicates greater measurement accuracy.

For the conventional method, the *R*^2^ was 0.921 (Fig. 2n, dashed line) and the RMSE was 24.9 (Fig. 2o, dashed line). For the proposed deep learning-based method, when the training image set size was *N* = 2, the values obtained did not match those of the ground truth, showing a substantially lower *R*^2^ and higher RMSE values (Fig. 2b, n, o). However, when the number of training images was *N* = 5 or more, the deep learning-based method consistently demonstrated equivalent or greater accuracy with higher *R*^2^ and lower RMSE values compared with the conventional method (Fig. 2c–o). In the case where *N* = 80, the *R*^2^ value was consistent with that of the conventional method at 0.921, while the RMSE value of 26.2 was higher than that of the conventional method, possibly because the properties of the additional training data differed substantially from those of the test data. Except for this instance, the proposed method showed superior accuracy than the conventional method.

Parallelness is an indicator of the alignment of the cytoskeletons; it reaches its maximum value of 1 when all cytoskeletons are oriented in the same direction, and approaches its minimum value of 0 as the orientations diverge (Higaki 2017). The measurements of parallelness, in contrast to the angle, showed weaker agreement with the ground truth for both the conventional and proposed methods, but the trend was similar. Specifically, for training sets of *N* = 5 or more, the proposed method exhibited equivalent or superior accuracy compared with the conventional method, which showed an *R*^2^ value of 0.932 and an RMSE of 0.143 (Fig. 3).

**Figure 3.**
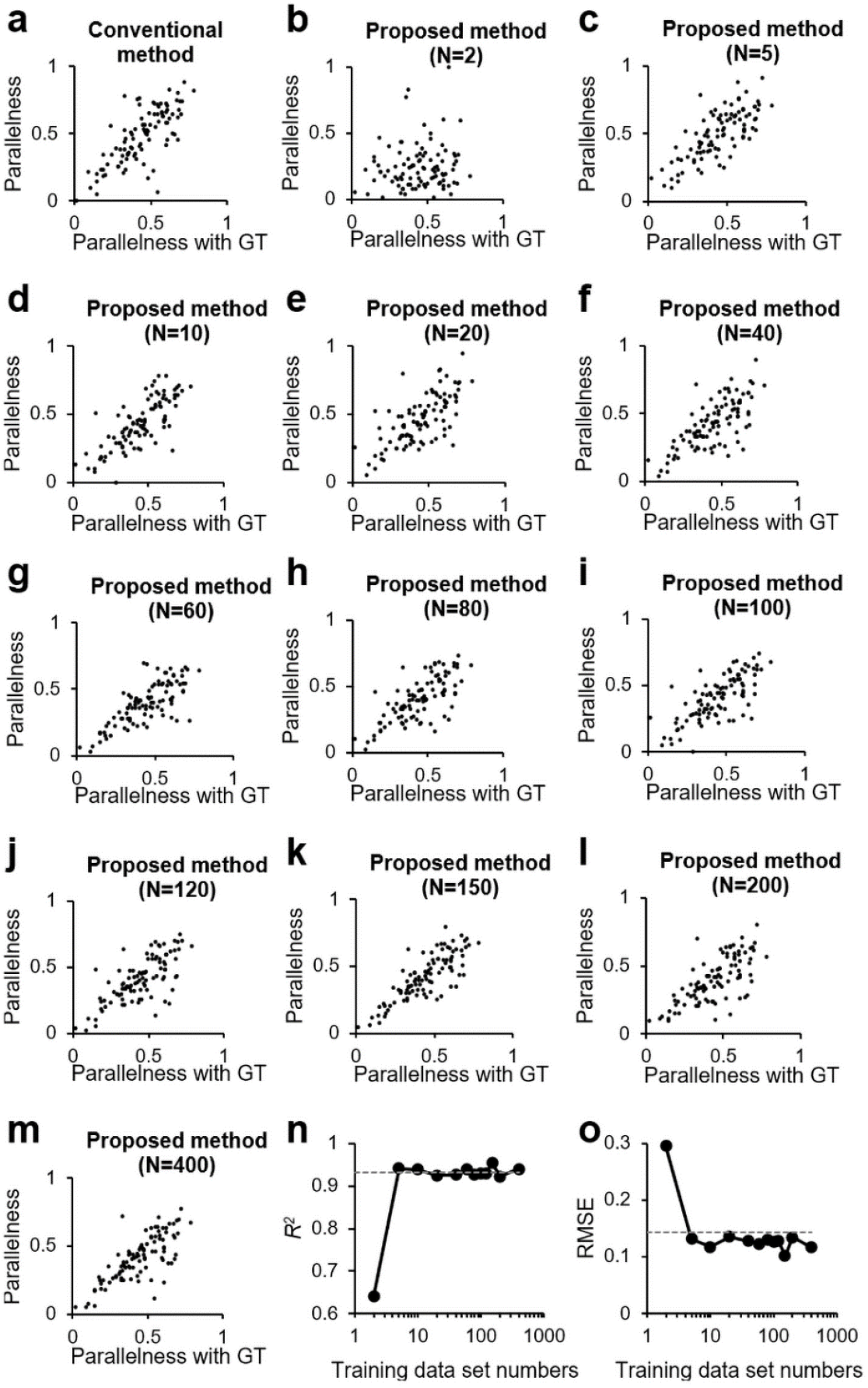
Evaluation of image processing pipelines for measurement of cytoskeleton parallelness. (a) Scatterplot comparing cytoskeleton parallelness values measured by the ground truth (GT) against those obtained via the conventional method. (b–m) Scatterplots comparing GT cytoskeleton parallelness values to those measured by the proposed method, varying the number of training image datasets at *N* = 2 (b), 5 (c), 10 (d), 20 (e), 40 (f), 60 (g), 80 (h), 100 (i), 120 (j), 150 (k), 200 (l), and 400 (m). (n, o) Plot of *R*^2^ (n) and root mean squared error (RMSE) (o) as an accuracy metrics, shown against the number of training datasets in the proposed method. The dashed line represents the value of 0.932 and 0.143 for the conventional method, respectively.

Density is the total length of skeletonized cytoskeleton per unit area (Higaki 2017). Therefore, it was expected to be the most demanding measure of segmentation accuracy. Indeed, for density, the trend differed from those for angle and parallelness. With the proposed method, up to a training set size of *N* = 20, the accuracy was slightly inferior to the conventional method, which showed an *R*^2^ value of 0.950 and an RMSE value of 1.52 (Fig. 4a–e, n, o). However, when the number of training images was greater than *N* = 100, the RMSE was lower than the value of the conventional method, confirming a stable improvement in accuracy (Fig. 4j–o). The *R*^2^ values for *N* = 120 (Fig. 4j), 150 (Fig. 4k), 200 (Fig. 4l), and *N* = 400 (Fig. 4m) were 0.967, 0.983, 0.963, and 0.978, respectively (Fig. 4j–n). The RMSE values for *N* = 120 (Fig. 4j), 150 (Fig. 4k), 200 (Fig. 4l), and *N* = 400 (Fig. 4m) were 1.17, 0.858, 1.26, and 0.988, respectively (Fig. 4j–m, o).

**Figure 4.**
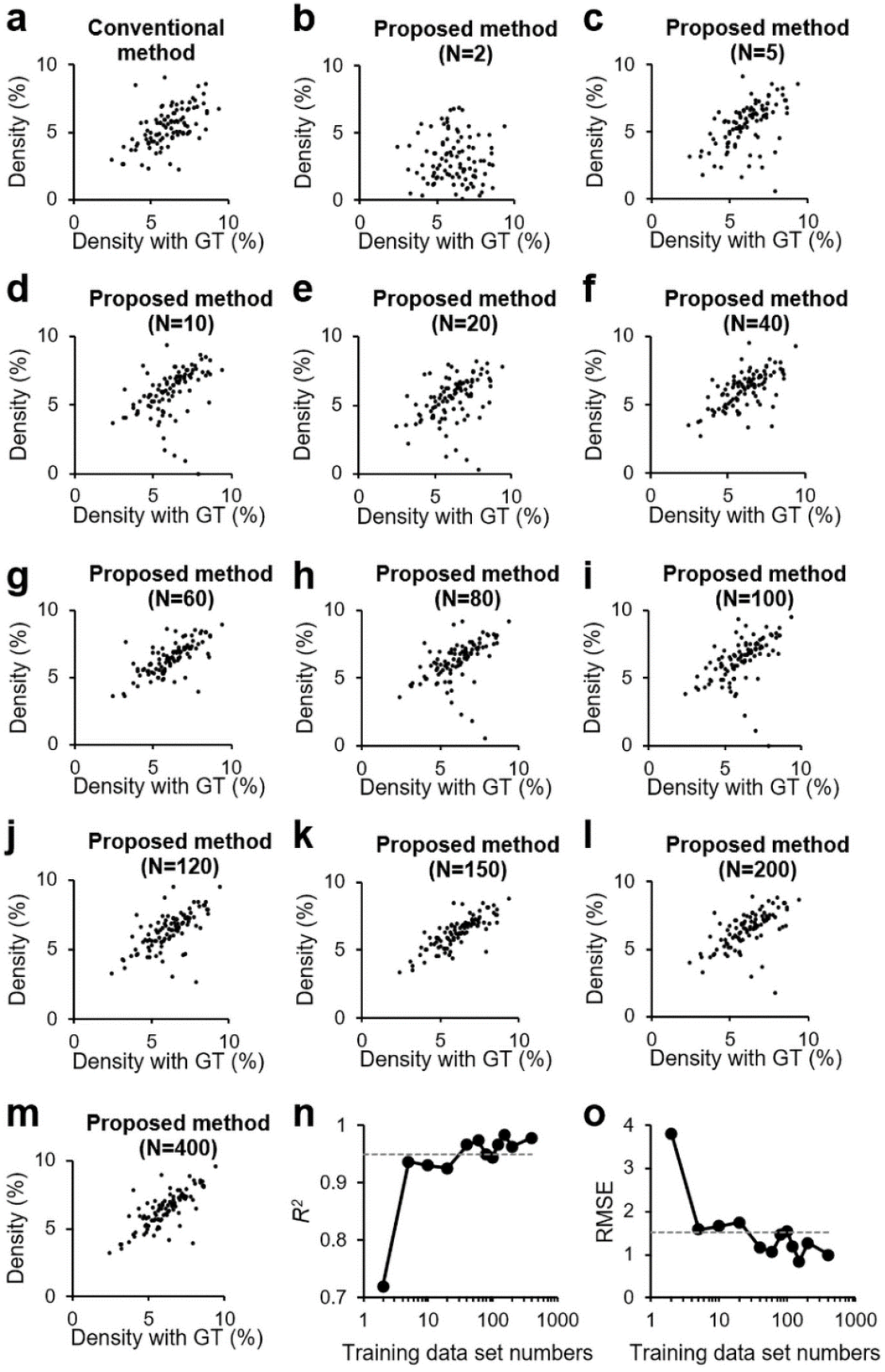
Evaluation of image processing pipelines for measurement of cytoskeleton density. (a) Scatterplot comparing cytoskeleton density values measured by the ground truth (GT) against those obtained via the conventional method. (b–m) Scatterplots comparing GT cytoskeleton density values to those measured by the proposed method, varying the number of training image datasets at *N* = 2 (b), 5 (c), 10 (d), 20 (e), 40 (f), 60 (g), 80 (h), 100 (i), 120 (j), 150 (k), 200 (l), and 400 (m). (n, o) Plot of *R*^2^ (n) and root mean squared error (RMSE) (o) as an accuracy metrics, shown against the number of training datasets in the proposed method. The dashed line represents the value of 0.950 and 1.52 for the conventional method, respectively.

These results suggested that, at least for the image sets used in this study, measurements of angle and parallelness were sufficiently accurate with the conventional methods or deep learning models trained on a limited number of images. However, the density measurements highlighted significant differences between the methodologies. The conventional method consistently failed to yield accurate results, revealing its limitations. Conversely, the proposed method, employing a deep learning model trained with a sufficient number of images, facilitated automatic measurements that closely approximated the manually thresholded ground-truth accuracy. This efficacy supported the proposed method’s capability to perform precise segmentation of cytoskeletons, leading to accurate and reliable automatic density measurements.

### Versatility of the proposed method for measuring cytoskeleton density in guard cells

To validate the versatility of the proposed method in cytoskeleton density measurements, we explored its utility for analysis of cytoskeletons in *A. thaliana* stomatal guard cells. This application was selected because the cytoskeleton density of the guard cells is associated with changes in the stomatal aperture (Higaki et al. 2010). The image processing pipeline employed was essentially identical to that used for BY-2 cells, but included additional steps to enhance cytoskeleton structures and specifically target guard cell regions (Supplementary Figure S1). For training of the deep learning segmentation model, we used 50 confocal images of actin filaments labeled with GFP-mTn, available from the LIPS image database (Higaki et al. 2012, 2013). In addition, we performed density measurements on 50 confocal images of actin filaments labeled with GFP-ABD2 in guard cells, newly captured for this study, and 50 confocal images of microtubules labeled with GFP-tubulin in guard cells, also sourced from the LIPS database. The results demonstrated that the *R*^2^ values were consistently higher for the proposed model, and the RMSE values were lower for both actin and microtubule images, indicating improved accuracy (Fig. 5a, b). Scatter plots revealed that the conventional method tended to underestimate density when compared with the ground truth (Fig. 5a, b). These findings suggested that the proposed method provides more accurate segmentation and density measurements in guard cells than the conventional method (Fig. 5a, b).

**Figure 5.**
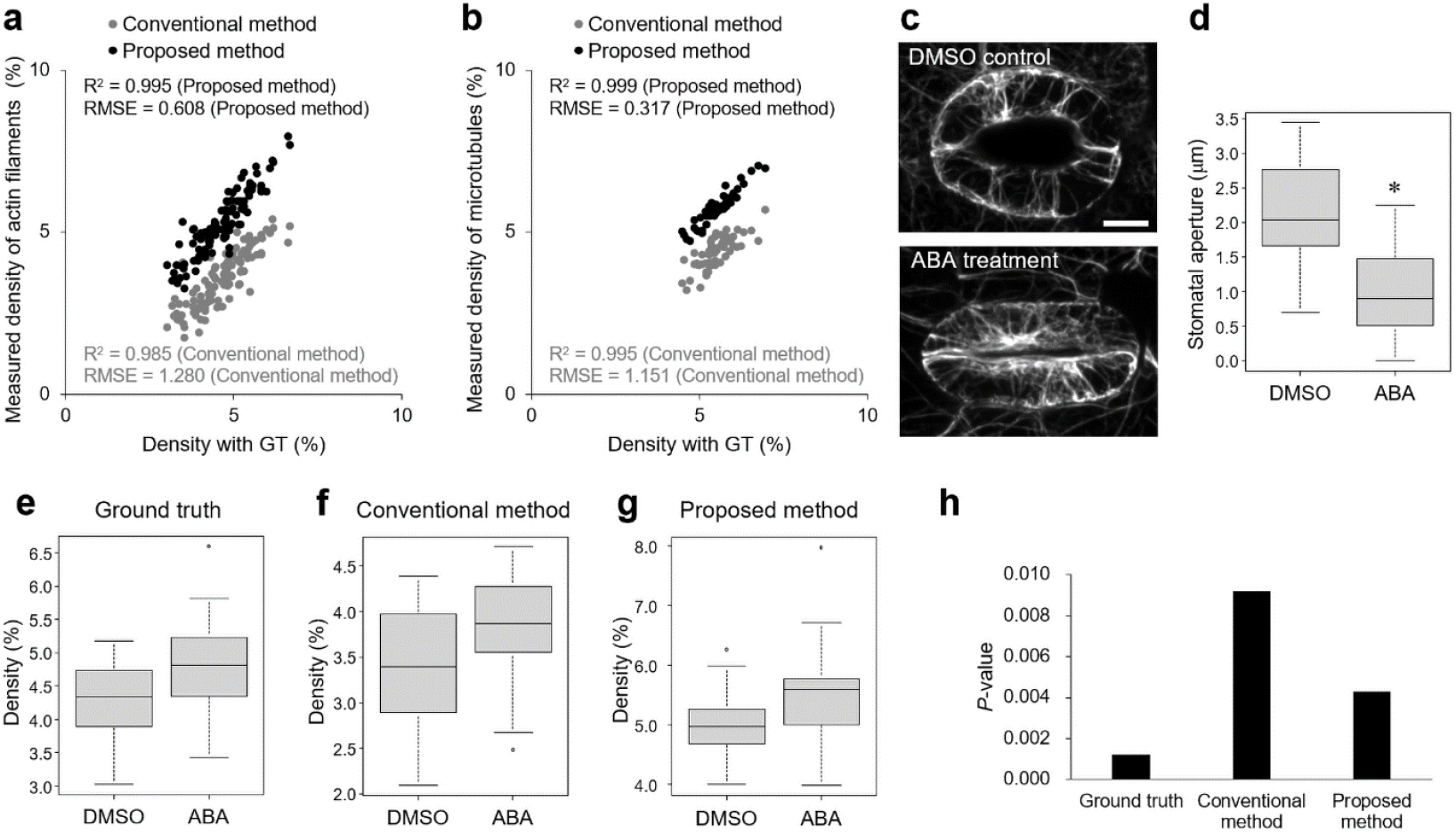
Accuracy evaluation of the proposed cytoskeleton segmentation in *Arabidopsis thaliana* guard cell images. (a, b) Scatterplots of cytoskeleton density values measured by the ground truth (GT) against those determined by the conventional method (gray points) and the proposed method (black points) for GFP-ABD2-labelled actin filaments (a) and GFP-tubulin (b). The *R*^2^ and root mean squared error (RMSE) are provided as accuracy metrics. The deep learning model was trained with the image dataset of GFP-mTn-labeled actin filaments in guard cells. (c) Confocal images of GFP-ABD2-labeled actin filaments in *A. thaliana* guard cells, treated with DMSO (control) or abscisic acid (ABA) for 5 min. Scale bar represents 10 μm. (d) Boxplot of stomatal aperture in guard cells expressing GFP-ABD2 treated with DMSO (control) or ABA for 5 min. An asterisk indicates statistical significance (*P* < 0.01, *U*-test, *N* = 30). (e–g) Boxplots of actin filament density measured by the ground truth (e), conventional (f), and proposed methods (g) under the DMSO (control) and ABA treatments. (h) *P*-values from *U*-tests comparing the actin filament density measurements across the ground truth (e), conventional (f), and proposed methods (g) under the DMSO (control) and ABA treatments.

To evaluate whether the proposed method could efficiently and accurately determine the physiological responses of actin filaments in guard cells, we analyzed images of *A. thaliana* cotyledons stably expressing GFP-ABD2 that were treated with ABA to induce stomatal closure. Actin filament density increases during ABA-induced stomatal closure (Shi et al. 2022). Upon treatment with 10 μM ABA, stomatal closure was observed (Fig. 5c, d). Density measurements of actin filaments in the DMSO control and ABA-treated samples were performed using manual thresholding for the ground truth (Fig. 5e), the conventional method (Fig. 5f), and the proposed method (Fig. 5g), respectively. The density of actin filaments was higher in ABA-treated samples for all methods (Fig. 5e–g). Comparative analysis using the *U*-test revealed *P*-values of 0.00122, 0.00918, and 0.004313, respectively, indicating that the proposed method provided significantly more precise segmentation and density measurements (Fig. 5h).

### Versatility of the proposed method for measuring cytoskeleton density in zygotes

To further explore the adaptability of the proposed method, we applied it to measure cytoskeleton density in *A. thaliana* zygotes using two-photon excitation microscopy. *Arabidopsis thaliana* zygotes were selected due to documented microtubule polarization towards the apical region during zygote elongation, which illustrates their potential for assessment of the broader applicability of the proposed technique (Kimata et al. 2016; Hiromoto et al. 2023). We captured 30 images of elongating zygotes, dividing them into two groups: 15 images for training the deep learning segmentation model and 15 images for validating its accuracy. The image processing pipelines were identical to that used for the guard cells. To specifically analyze microtubule polarization, we designated regions of interest of 35 pixels × 35 pixels in each of the apical and basal regions of the zygotes and assessed the microtubule density within these regions (Fig. 6a, yellow squares). A higher microtubule density was observed in the apical region compared with the basal region for all methods (Fig. 6b). Comparative scatter plots, *R*^*2*^ and RMSE analysis against the ground truth demonstrated that the proposed method provided more precise measurements than the conventional approach (Fig. 6c). In addition, we quantified microtubule polarization by comparing the ratio of apical to basal microtubule densities (Fig. 6d). The conventional method occasionally produced unstable results, overestimating this ratio, indicative of segmentation inaccuracies in the basal region, which presumably resulted in an artificially low denominator, leading to inflated ratios (Fig. 6d). These findings further demonstrated the utility of the proposed method for accurate quantification of microtubule density in zygotes, confirming its broader applicability to different cellular contexts.

**Figure 6.**
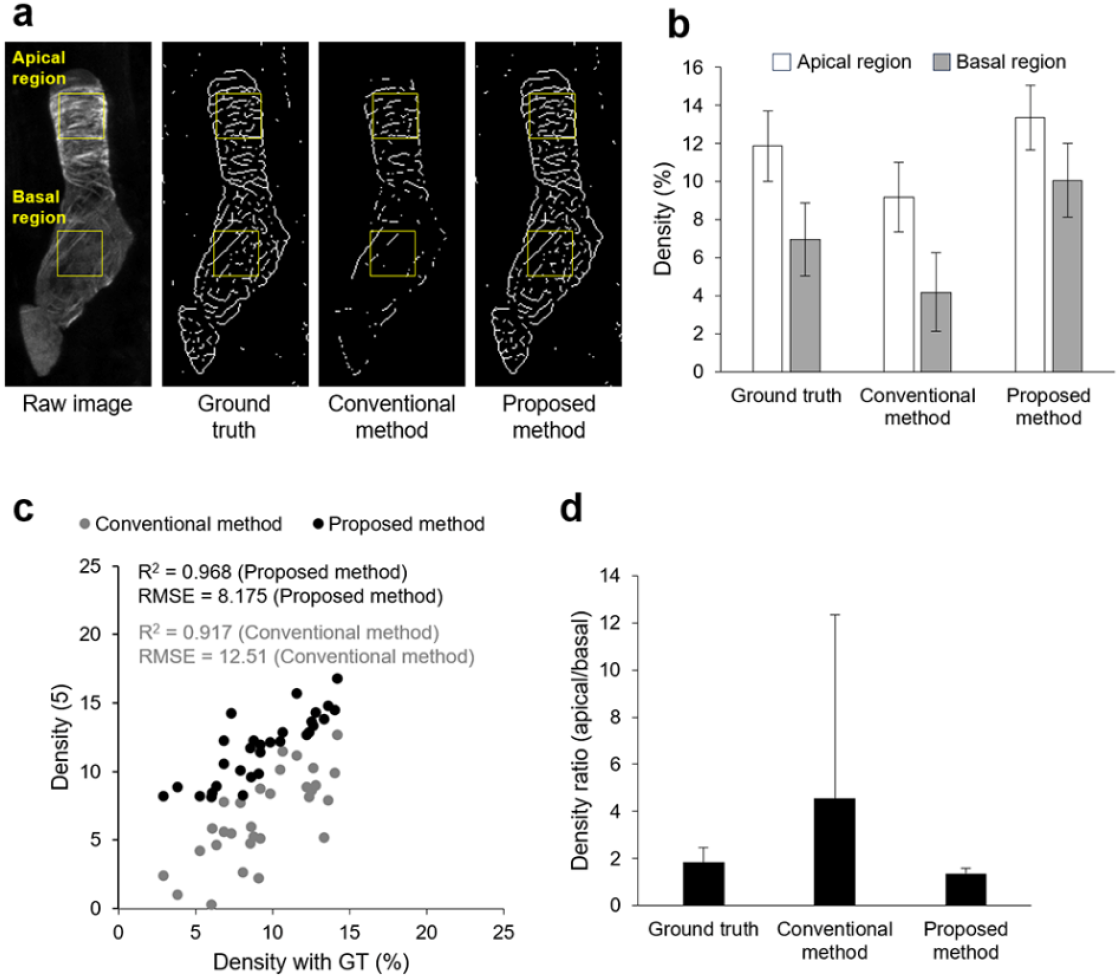
Accuracy evaluation of the proposed cytoskeleton segmentation using two-photon excitation microscopic images of *Arabidopsis thaliana* zygotes. (a) Representative input (raw) images and corresponding skeletonized outputs (ground truth, conventional, and proposed methods) showing microtubules in *A. thaliana* elongating zygotes expressing Clover-TUA6. Yellow squares indicate the apical and basal regions. The width of the square is 7.11 μm. (b) Quantification of microtubule density in the zygote apical (white column) and basal regions (gray column) as measured by the ground truth, conventional, and proposed methods. Data are the mean of *N* = 15; error bars indicate the standard deviation. (c) Scatter plot comparing microtubule density values of the apical and basal regions ascertained by the ground truth (GT) with those derived from the conventional (gray points) and proposed methods (black points) (*N* = 30). The *R*^2^ and root mean squared error (RMSE) are provided as accuracy metrics. (d) Ratio of microtubule density in the apical region relative to that in the basal regions. Data are the mean of *N* = 15; error bars indicate the standard deviation. Note that values obtained by the conventional method were potentially overestimated because of underdetection of basal microtubules.

## Discussion

In this study, we systematically evaluated the utility of deep learning-based methods for cytoskeleton segmentation compared with conventional methods that employ automated thresholding algorithms for analysis of fluorescence microscopic images of actual plant cytoskeletons. Our assessment using confocal microscopic images of cortical microtubules in tobacco BY-2 cells revealed that the deep learning-based segmentation method achieved high precision, especially for measurement of cytoskeleton density (Fig. 4). Conversely, conventional methods proved sufficient for assessment of angle and parallelness (Figs. 2 and 3). These results indicated that, even if segmentation accuracy was compromised, the correct identification of principal cytoskeleton structures allowed for reliable measurement of their orientation and parallelness. This applicability was expected to extend across various biological contexts, although exceptions may occur under specific conditions where cytoskeletons at low intensities display unique orientations, which is unlikely to be assumed in practice.

As anticipated, our systematic validation with actual microscopic images confirmed the critical importance of segmentation accuracy for reliable cytoskeleton density measurements in confocal microscopic images of tobacco BY-2 cells (Fig. 4), *A. thaliana* guard cells (Fig. 5), and two-photon excitation microscopic images of *A. thaliana* zygotes (Fig. 6). Automated threshold determination algorithms, such as Otsu’s method, while invaluable as image processing tools, must be employed with caution when precise measurements are essential. The deep learning-based segmentation method evaluated in this study effectively automated the threshold determination process while maintaining accuracy in cytoskeleton segmentation. This feature would be particularly beneficial for advanced imaging applications, such as long-term time-lapse imaging or extensive analysis of numerous samples captured with automated confocal microscopy, where the volume of image data is substantial (Cui et al. 2023).

However, deep learning-based segmentation has certain limitations at present. The model used in this study heavily relies on the specificity of its training data, thus requiring the creation of new datasets for different cell types or conditions. Notably, while the model trained with images of GFP-mTn-labeled actin filaments in guard cells effectively analyzed images of GFP-ABD2-labeled actin filaments and microtubules in the same cell types (Fig. 5), it showed restricted performance when applied to other cell types, such as guard cells and zygotes, when trained on images from BY-2 cells. This underlines the critical need to enhance the adaptability of the model across diverse biological contexts. Future research should focus on developing versatile deep learning models that can accommodate a variety of cell types and conditions. This method may also be applicable to the analysis of other fibrous structures, such as cellulose microfibers, as well as the cytoskeleton (Kuki et al. 2017). To enhance the generalizability of deep learning models, it is generally essential to expand the training image datasets to include a diverse variety of cell types and conditions. Initially, it is crucial to collect images from different plant species and cell types, incorporating various fluorescent probes, staining methods, and stages of cellular development. Furthermore, datasets should not only include images from confocal and two-photon excitation microscopy, but also from other modalities, such as light-sheet microscopy (Ovečka et al. 2022), variable-angle epifluorescence microscopy (Konopka and Bednarek 2008; Higaki 2015), and super-resolution microscopy (Komis et al. 2015; Ovečka et al. 2022). This integration will contribute to improving the model’s versatility. In addition, data augmentation techniques can be employed to generate a larger set of training data from existing images (Chlap et al. 2021). By applying transformations, such as rotation, flipping, scaling, and adding noise, we can train the model to be robust under diverse imaging conditions. These strategies will enable deep learning models to be applicable to a broader range of plant materials and research questions, not only enhancing the efficiency of experiments but also providing novel biological insights in the field of plant cell biology.

Several software tools for quantitatively evaluating cytoskeleton organization are available in addition to our technique. In the field of plant science, tools such as Fibril Tool (Boudaoud et al. 2014), OrientationJ (Püspöki et al. 2016), and MicroFilament Analyzer (Jacques et al. 2013) are widely used. In these tools, segmentation is not always necessary when evaluating orientation quantitatively, for example. In this study, we proposed a method that combines accuracy and automation in threshold setting for cytoskeleton segmentation using deep learning. However, deep learning-based image transformation may not only aid in segmentation but also be applicable to these established tools. In other words, the deep learning output images, which are enhanced representations of the cytoskeleton prior to binarization, could be used as input for existing tools, potentially improving quantitative evaluations. Although this remains a subject for future investigation, deep learning-based filtering may also be valuable for quantitative evaluations that do not require segmentation.

In conclusion, the present findings validated the superiority of deep learning in improving cytoskeleton segmentation and measurement of cytoskeleton density, indicating its potential to revolutionize quantitative microscopy in plant cell biology. Continued development and refinement of these approaches are expected to expand their utility across a broader range of cell types and conditions, thereby enhancing both the efficiency of experiments and the depth of biological insights.

## Acknowledgements

This work was supported by the Japan Science and Technology Agency (CREST; JPMJCR2121) to Takumi Higaki and Minako Ueda, and by the Japan Society for the Promotion of Science [Advanced Bioimaging Support (JP22H04926 to Minako Ueda); a Grant-in-Aid for Early-Career Scientists (JP22K15135 to Hikari Matsumoto); and a Grant-in-Aid for Scientific Research (B) (JP19H03243 to Minako Ueda)]. We thank Ms. Hitomi Okada (Kumamoto University) and Ms. Remi Kawakami (Kumamoto University) for their assistance with plant maintenance. We thank Robert McKenzie, PhD, from Edanz (https://jp.edanz.com/ac) for editing a draft of this manuscript.

## Supplementary Figure

**Supplementary Figure S1.**
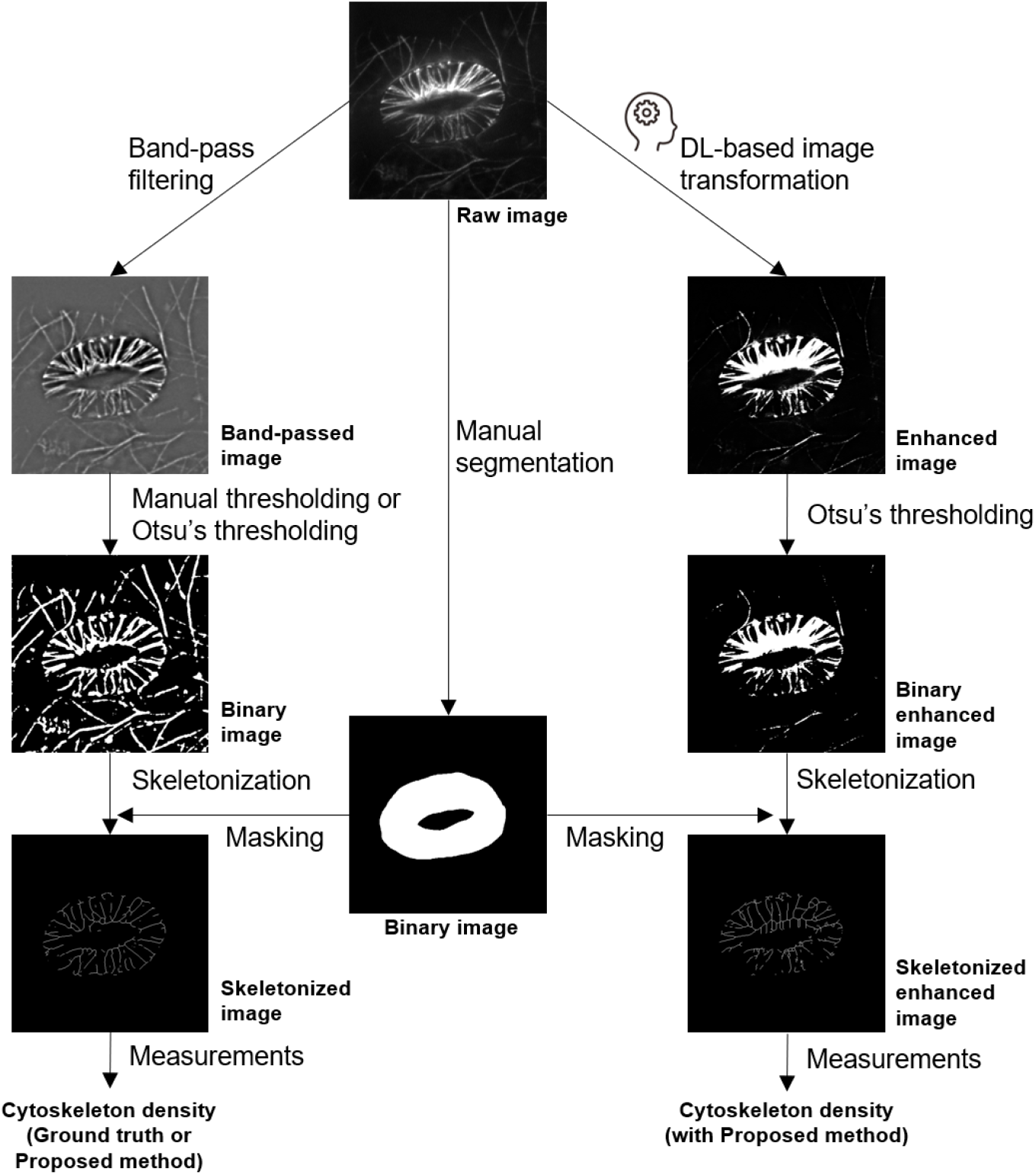
Image processing pipelines for evaluation of segmentation accuracy in *Arabidopsis* guard cell images. The workflow was essentially identical to that used for tobacco BY-2 cells, as shown in Figure 1, with the addition of a band-pass filtering and masking procedure.

